# The olfactory ventricle: an overlooked and evolutionarily conserved feature of the mammalian brain

**DOI:** 10.64898/2026.10.04.756487

**Authors:** Lucile Butruille, Jean-Philippe Dubois, Martine Migaud, Sylvie Remaud

## Abstract

The olfactory ventricle (OV), an anterior extension of the lateral ventricular system into the olfactory bulbs, is generally considered a transient developmental structure that closes during early postnatal life. However, this view largely derives from a limited number of laboratory models, and the evolutionary distribution and cellular organization of the adult OV remain poorly understood. Here, we combined a comparative survey of mammalian neuroanatomy with phylogenetic analyses and cellular characterization of the OV across species. We compiled anatomical data for 154 mammalian species spanning 26 orders and 80 families, revealing widespread persistence of a patent OV into adulthood. Phylogenetic analyses supported OV presence as the ancestral mammalian state and indicated that its absence is phylogenetically structured and consistent with repeated secondary losses. Chiroptera displayed multiple lineage-specific losses, whereas variation within Muridae extended to the intraspecific level, with contrasting OV persistence among *Mus musculus* and *Rattus norvegicus* populations. Comparative histological analyses in sheep, *Peromyscus maniculatus*, and CD1 mice revealed that, despite marked interspecific differences in OV morphology, the persistent ventricular region retains a conserved cellular organization characterized by progenitor- and glial-associated markers, proliferative cells, immature neuronal populations, and a differentiated apical/ciliary interface. Although these observations do not establish local neuronal production or demonstrate that the OV constitutes an autonomous neurogenic niche, their convergence across species indicates that the persistent adult OV is not simply an anatomically residual lumen but a specialized component of mammalian olfactory bulb organization. Together, our findings identify the OV as an ancestral and widely conserved feature of the mammalian brain and provide an evolutionary and cellular framework for investigating the developmental mechanisms controlling its persistence and its potential contribution to adult olfactory neurogenesis and plasticity.

## 1. Introduction

Adult olfactory neurogenesis is one of the best-characterized examples of persistent neuronal production in the mammalian brain (Lim & Alvarez-Buylla, 2016). Much of our current understanding derives from laboratory mice, particularly the C57BL/6 strain, in which NSPCs located in the subventricular zone (SVZ) generate neuroblasts that migrate along the rostral migratory stream (RMS) before reaching the olfactory bulb (OB), where they differentiate into interneurons. This SVZ-RMS-OB axis has consequently become the canonical representation of adult olfactory neurogenesis (Doetsch et al., 1999; Lim & Alvarez-Buylla, 2016; Ming & Song, 2011).

Yet, this anatomical and neurogenic organization is not established from the outset of brain development. During embryonic development, OB progenitors are located in the lateral ganglionic eminence and its rostral extension, the olfactory ventricular zone, which lines the olfactory ventricle (OV) and contributes progenitors to neuronal production within the developing OB. During the early postnatal period, the OV progressively collapses, ultimately leaving a collapsed ventricular remnant associated with the RMS (Chen et al., 2017; Jacquet et al., 2011). The embryonic ventricular organization is thereby transformed into the postnatal SVZ-RMS system, which subsequently supports the long-distance migration of newly generated neuroblasts toward the OB. Thus, the classical SVZ-RMS-OB axis of the adult laboratory mouse represents the outcome of a profound developmental remodeling of an initially ventricular neurogenic system rather than an anatomical organization that is fixed throughout development.

This developmental trajectory raises a fundamental comparative question: is closure of the OV, as described in the laboratory mouse, representative of the mammalian condition? Several observations suggest that it may not be. Comparative analyses of the adult SVZ have highlighted substantial interspecific diversity in its organization. In particular, whereas the OV closes during development in commonly studied laboratory rodents, an open OV has been reported in the adult OB of several other mammals, including rabbits (Luzzati et al., 2003), sheep (Brus et al., 2013), opossums (Tepper et al., 2021), wild dogs (Chengetanai et al., 2020) and some non-human primates, such as macaques, rhesus monkeys, and common marmosets (Gil-Perotin et al., 2009; Sawamoto et al., 2011; Wang et al., 2011). The situation in humans further illustrates the complexity of OV persistence. Although developmental studies have traditionally suggested that the olfactory ventricular cavity becomes obliterated before birth (Smitka et al., 2009) reported OV-like structures by magnetic resonance imaging in 72 of 122 healthy adults (59%), including bilateral structures in 43% of individuals. Their frequency was higher in younger than in older adults, suggesting possible variation in OV persistence throughout life, although the ventricular nature of such MRI signals has subsequently been debated. Together, these observations challenge the assumption that the closed configuration familiar from laboratory mice represents the general mammalian condition.

Despite these observations, the distribution of the OV has never been examined systematically across mammalian diversity. Evidence for its presence is scattered throughout more than a century of literature, including comparative neuroanatomical descriptions, histological studies, brain atlases and, more recently, neuroimaging studies. Consequently, the evolutionary history of the OV remains largely unresolved. Which mammalian lineages retain an open OV into adulthood, and in which lineages is it lost through developmental closure? Addressing this question requires moving beyond a small number of classical laboratory models and examining olfactory ventricular anatomy across the breadth of mammalian diversity.

Beyond its evolutionary significance, persistence of the OV may also have important implications for the organization of adult olfactory neurogenesis. During mouse embryonic development, the OV is bordered by a proliferative ventricular zone that contributes progenitors to OB neurogenesis, whereas its early postnatal collapse accompanies the transition toward the SVZ-RMS-OB organization (Jacquet et al., 2011). If an open OV and its associated ventricular environment persist into adulthood in other mammals, features of this local neurogenic organization might therefore also be retained. Adult olfactory neurogenesis has previously been documented in sheep. Proliferating BrdU- and Ki67-positive cells have been identified within the adult main OB (Brus et al., 2010), and subsequent work demonstrated the contribution of SVZ-derived precursors to newly generated olfactory cells in this species (Brus et al., 2013). These observations establish that the adult sheep OB receives newly generated cells, but do not resolve whether part of this population could also originate locally from progenitors associated with the OV. These observations raise the hypothesis that persistence of an open OV may be associated with a ventricular wall retaining cellular features compatible with a local neurogenic niche within the adult OB, potentially complementing neuronal recruitment through the classical SVZ-RMS pathway.

Here, we reconstruct the evolutionary history of the OV across mammals through an extensive survey of the available literature, integrating evidence from comparative neuroanatomy, histology, brain atlases, magnetic resonance imaging and other anatomical sources. By sampling species across the major mammalian lineages, we characterize the diversity of olfactory ventricular anatomy and determine which lineages retain an open OV into adulthood and which undergo its developmental closure. We further examine the cellular organization of the OV in sheep, *Peromyscus maniculatus*, and two strains of *Mus musculus*, CD1 and C57BL/6, to determine whether persistence of the OV is associated with a ventricular organization compatible with a neurogenic niche and with the presence of cell populations involved in adult neurogenesis. Together, these approaches investigate both the evolutionary conservation of the OV and the possibility that its persistence may contribute to an alternative organization of adult olfactory neurogenesis across mammals.

## 2. Methods

### 2.1. Literature survey and species selection

To investigate the distribution of the OV across mammals, we conducted an extensive survey of published comparative neuroanatomical literature and publicly available neuroanatomical resources. Published sources included primary research articles, histological and neuroanatomical studies, anatomical and stereotaxic brain atlases, magnetic resonance imaging studies, serial brain sections, and historical neuroanatomical monographs. In addition, online comparative brain databases and digital atlases, including the Comparative Mammalian Brain Collections and BrainMaps, were examined when they provided histological sections allowing direct visualization of the internal anatomy of the OB. Literature searches combined scientific and common species names with terms related to the OB and ventricular system, including OV, OB ventricle, olfactory ventricular cavity, olfactory vesicle, rhinocele, ventricle, and ventricular system. Reference lists of relevant publications were additionally screened to identify older anatomical sources not retrieved through electronic searches. Species were included only when the available material allowed direct assessment of the internal anatomy of the OB. Adult specimens were prioritized, as the primary objective was to determine whether an open OV persists into adulthood. Taxonomic information was cross-checked against current mammalian taxonomic databases, whereas anatomical information used to determine OV status was derived exclusively from the original neuroanatomical sources or directly examined histological and imaging resources described above.

### 2.2. Classification of OV anatomy

For each species, OV status was classified as present or absent based on direct anatomical evidence. The OV was scored as present when a clearly identifiable open ventricular cavity was observed within the OB of an adult individual. It was scored as absent when the available anatomical material provided adequate visualization of the internal organization of the adult OB but no ventricular cavity was observed. Importantly, “absent” therefore refers specifically to the absence of a patent OV in the adult OB and does not imply that an OV was absent earlier in development.

When several sources were available for the same species, all informative observations were considered, with priority given to material providing the clearest visualization of the internal anatomy of the OB. Histological sections and other sources allowing direct visualization of the ventricular cavity were favored over indirect anatomical descriptions. Particular attention was paid to section orientation, tissue preservation, and image resolution, as these factors may influence the detection of small ventricular cavities.

### 2.3. Phylogenetic analysis / Ancestral state reconstruction

Species names were taxonomically reconciled with the mammalian phylogenies of Upham et al. (2019). When current taxonomic nomenclature differed from that used in the phylogeny, species were matched to the corresponding phylogenetic tip while retaining the current name in the comparative dataset. Because the comparative dataset included multiple observations for some species, as well as taxonomic entries corresponding to the same phylogenetic tip, the number of phylogenetic units was lower than the number of species-level records in the original survey. Concordant observations mapping to the same phylogenetic tip were collapsed into a single state. Following taxonomic reconciliation and consolidation of repeated observations, the dataset comprised 145 unique phylogenetic tips. Phylogenetic uncertainty was incorporated using 100 completed mammalian phylogenies derived from the phylogenetic framework of Upham et al. (Upham et al., 2019) and distributed through the megatrees resource. This set comprised 50 phylogenies based on the Node Dating Exponential backbone and 50 based on the Fossilized Birth-Death backbone. Each tree contained 5,911 mammalian species and was pruned to the taxa represented in our dataset prior to analysis. OV status was treated as a binary character, with the presence of an open OV coded as 1 and the absence of a patent OV in the adult OB coded as 0.

Ancestral-state reconstruction was performed in Python using a maximum-likelihood implementation of a two-state continuous-time Markov model. Two models of character evolution were fitted independently to each of the 100 phylogenies: an equal-rates model (ER), in which transitions between OV presence and absence were constrained to occur at the same rate, and an all-rates-different model (ARD), in which the transition rates from absence to presence and from presence to absence were estimated independently. Stationary frequencies implied by the estimated transition-rate matrix were used as root-state priors. Model fit was compared using Akaike’s Information Criterion (AIC). For each phylogeny, marginal probabilities of OV absence and presence at the root were calculated under the ARD model. Analyses were implemented in Python 3.13.5 using NumPy 2.3.5 for numerical operations, SciPy 1.17.0 for maximum-likelihood optimization, and Biopython 1.86 for phylogenetic tree manipulation. Transition-rate parameters were optimized on the log scale by numerical maximization of the phylogenetic likelihood using Felsenstein’s pruning algorithm (Felsenstein, 1981).

*Mus musculus* and *Rattus norvegicus* displayed intraspecific variation in OV status among the anatomical observations included in the survey. Both species were coded as OV-absent in the primary analysis. To determine whether this coding affected inference of the ancestral mammalian state, the analyses were repeated with both species alternatively coded as OV-present.

Finally, following probabilistic inference of the ancestral mammalian state, maximum-parsimony reconstruction was used to identify the minimum number and phylogenetic distribution of transitions required to explain the observed distribution. The root was constrained to OV-present, consistent with the probabilistic ancestral-state reconstruction. Transitions between OV presence and absence were assigned equal unit costs, and minimum-transition scenarios were evaluated across all 100 phylogenies. Particular attention was given to Chiroptera, in which OV absence occurred in several phylogenetically distinct lineages. Phylogenetic signal in OV distribution was quantified using the D statistic for binary traits following Fritz and Purvis (2010). For each of the 100 phylogenies, the observed sum of sister-clade differences was compared with distributions generated under two null models: random permutation of OV states among species and a Brownian threshold model preserving the observed prevalence of the binary character. Each null distribution was generated using 1,000 simulations. D values approaching 1 indicate a phylogenetically random distribution, whereas values approaching 0 indicate a level of phylogenetic clustering expected under a Brownian threshold model; negative values indicate stronger clustering. The analysis was performed using *M. musculus* and *R. norvegicus* coded as OV-absent and repeated with both species coded as OV-present as a sensitivity analysis.

### 2.4. Animals

Two months-old C57BL/6J and CD1 *Mus musculus* wild-type male mice were purchased from Janvier Labs (Le Genest-Saint-Isle, France). Two months-old wild-derived male deer mice (*Peromyscus maniculatus*) were obtained from the Peromyscus Genetic Stock Center (University of South Carolina, Columbia, SC, USA). Mice were housed in ventilated cages under a 12:12 h light-dark cycle for *Mus musculus* and 12:8 h light-dark cycle for *Peromyscus maniculatus*, with food and water available *ad libitum*.

Adult Ile de France ewes (2-years-old) were kept in sheepfold, under natural photoperiod. Animals were fed daily with alfalfa, corn, straw, and vitamin and mineral supplements, with free access to water.

### 2.5. Tissue preparation

Mice were deeply anesthetized with pentobarbital (130 mg/kg; Centravet) and intracardially perfused with 1X PBS followed by 4% paraformaldehyde (PFA) in 1X PBS. Brains were dissected, post-fixed overnight in 4% PFA in PBS at 4°C, and cryoprotected overnight in 30% sucrose in PBS at 4°C.

Ewes were decapitated, and the heads were perfused through the carotid arteries with 2 L of 1% sodium nitrite in 0.9% NaCl at 37°C, followed by 4 L of cold 4% PFA in 0.1 M phosphate buffer (pH 7.4). OB-containing tissue blocks were subsequently post-fixed for 48 h in the same fixative and cryoprotected in 20% sucrose at 4°C.

Before sectioning, mouse brains and sheep OBs were embedded in OCT compound (Sakura Finetek, The Netherlands) and stored at -80°C. Mouse OBs were cut coronally at 12 μm using a cryostat (Leica). Sections were directly mounted onto Superfrost Plus slides (Fisher Scientific, Illkirch, France) and stored at -80°C until immunohistochemical processing. Sheep OBs were cut coronally at 30 μm as free-floating sections and stored at -20°C in cryoprotectant solution until immunohistochemical processing.

### 2.6. Immunohistochemistry and imaging

For sheep OBs, free-floating sections were first mounted onto Superfrost Plus glass slides. Prior to immunostaining, mouse and sheep OB sections were rehydrated and immersed in citrate buffer for 10 min at room temperature. Sections were then subjected to heat-induced antigen retrieval in citrate buffer. Slides were placed in a water bath when the citrate buffer reached 60°C, and the temperature was progressively increased to 90°C. Sections were maintained at 90°C for 20 min. Heating was then stopped, and sections were allowed to cool progressively in the citrate buffer to room temperature for approximately 20-30 min. Slides were subsequently washed three times for 5 min in PBS. Sections were then incubated for 30 min at room temperature in blocking solution 1% BSA (Sigma), 0.3% Triton X-100 and 10% donkey serum (Sigma) in PBS. Sections were then incubated overnight at 4◦C with primary antibodies diluted in the same solution. Primary antibodies were: mouse anti-PCNA (1/500, Noovocastra), rabbit anti-MCM2 (1/500, Abcam), chicken anti-Vimentin (1/2000, Millipore), rabbit anti-SOX2 (1/400, Abcam), guinea pig anti-DCX (1/500, Sigma), chicken anti-GFAP (1/1200, Abcam), mouse anti-Tubulin acetylated (1/500, Sigma), mouse anti-β catenin (1/300 BD Transduction laboratories), mouse anti-CD133 (1/300, Invitrogen), ARL13b (1/700, NeuroMab), rabbit anti-AC3 (1/300, Santa Cruz). After 3 washes in PBS at room temperature, sections were incubated for 2 h with the appropriate Alexa Fluor-conjugated secondary antibodies (1% BSA, 0.3% Triton X-100 and 1% donkey serum in PBS). Sections were then coverslipped using Fluoromount-G containing DAPI (SouthernBiotech).

Fluorescence images were acquired using a Zeiss LSM 880 confocal microscope (Carl Zeiss Microscopy) using a 40X objective. Z-stacks were acquired from the OB region surrounding the OV. Images were processed using ImageJ.

## 3. Results

### 3.1. The OV is broadly distributed across mammalian lineages

Our comparative survey compiled 154 anatomical observations spanning 26 mammalian orders and 80 families (Table 1). The dataset encompassed the three major mammalian lineages, including two monotreme, 16 marsupial and 133 placental taxa, with representatives of Afrotheria, Xenarthra, Euarchontoglires and Laurasiatheria. All observations were obtained from adult specimens, and OV status was established from anatomical material allowing direct visualization of the internal organization of the OB, primarily histological sections and brain atlases, magnetic resonance imaging, and diceCT/µCT datasets.

**Table 1.**
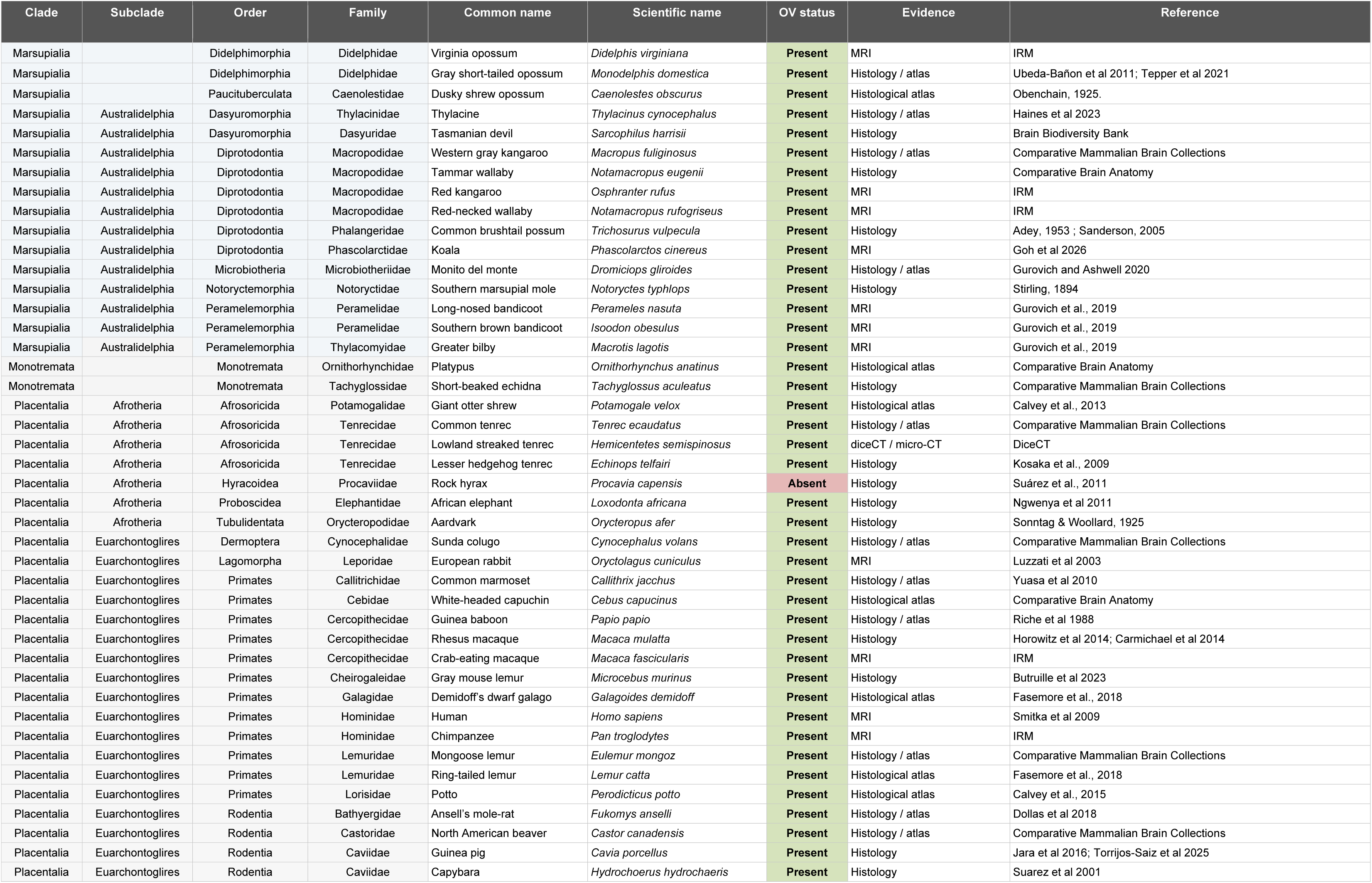
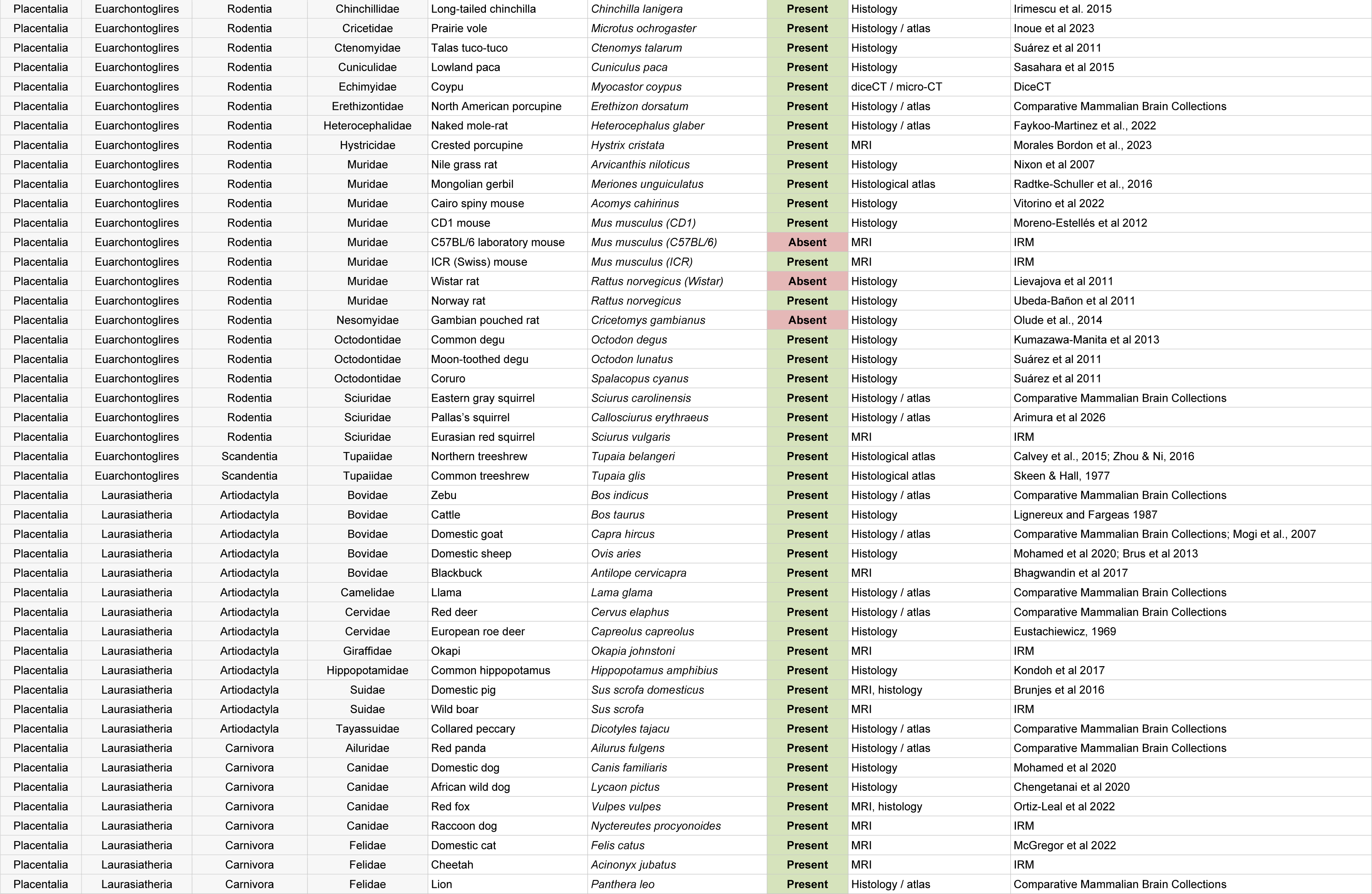
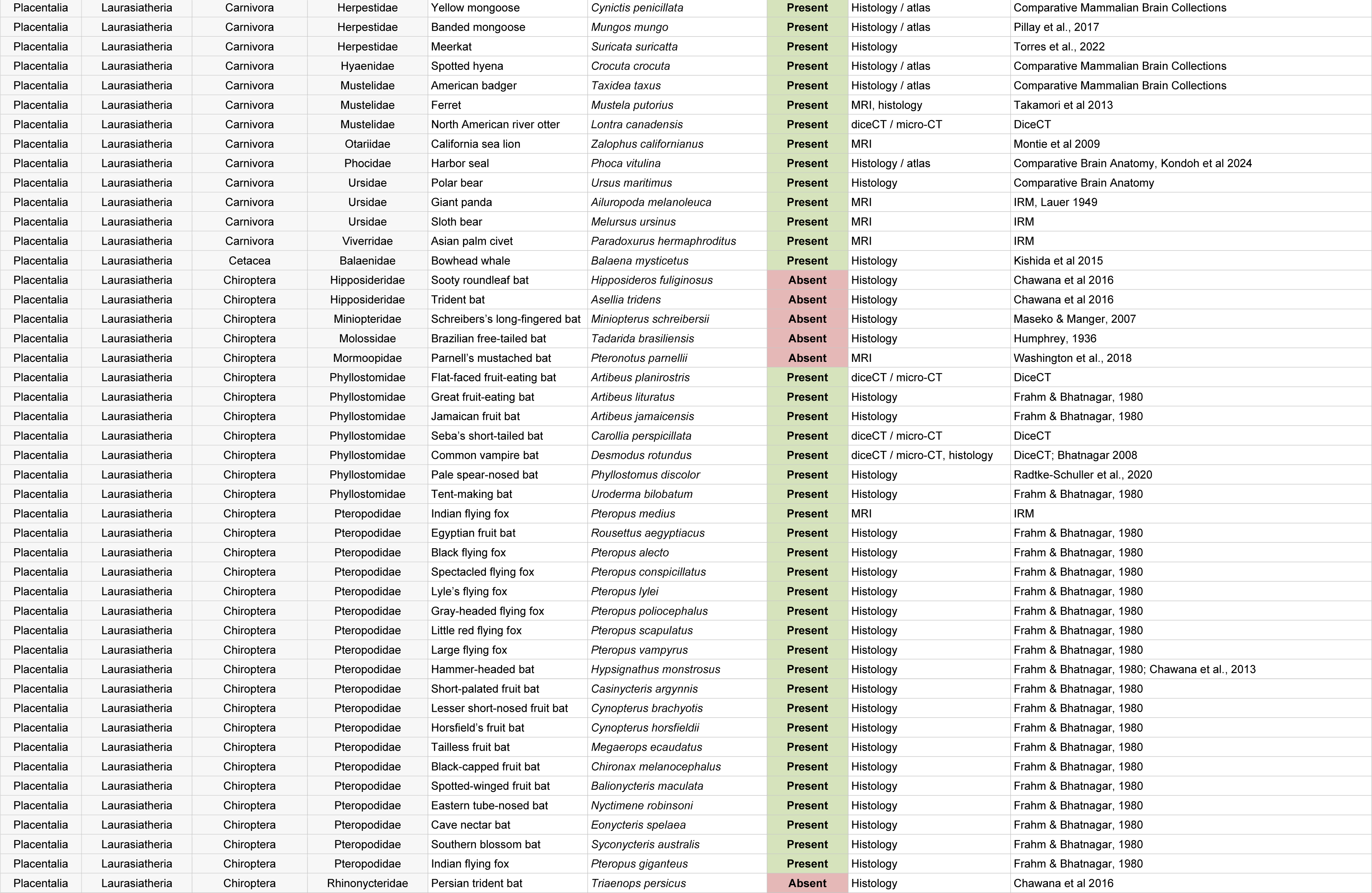
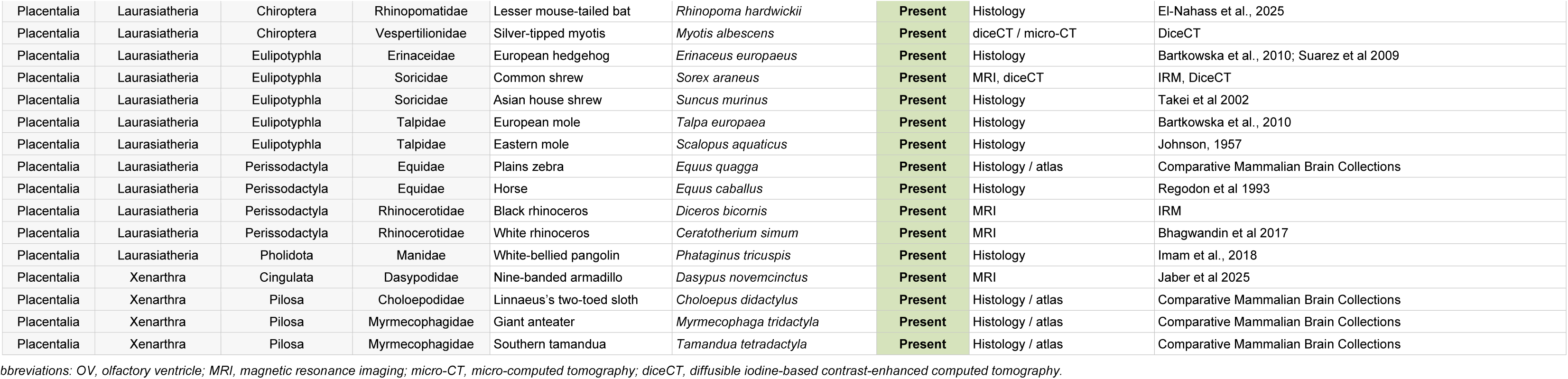
Comparative dataset of adult olfactory ventricle status across mammals. The table reports 151 adult anatomical observations spanning 26 mammalian orders and 80 families. OV status is based on direct anatomical evidence within the olfactory bulb.

An open OV was identified in 144 of the 154 observations, whereas 10 lacked a patent ventricular cavity within the adult OB. OV presence was documented throughout the mammalian phylogeny, including monotremes, marsupials and all major placental lineages represented in the dataset. In contrast, OV absence showed a markedly restricted taxonomic distribution, being detected only within Hyracoidea, Rodentia and Chiroptera (Fig. 1).

**Figure 1.**
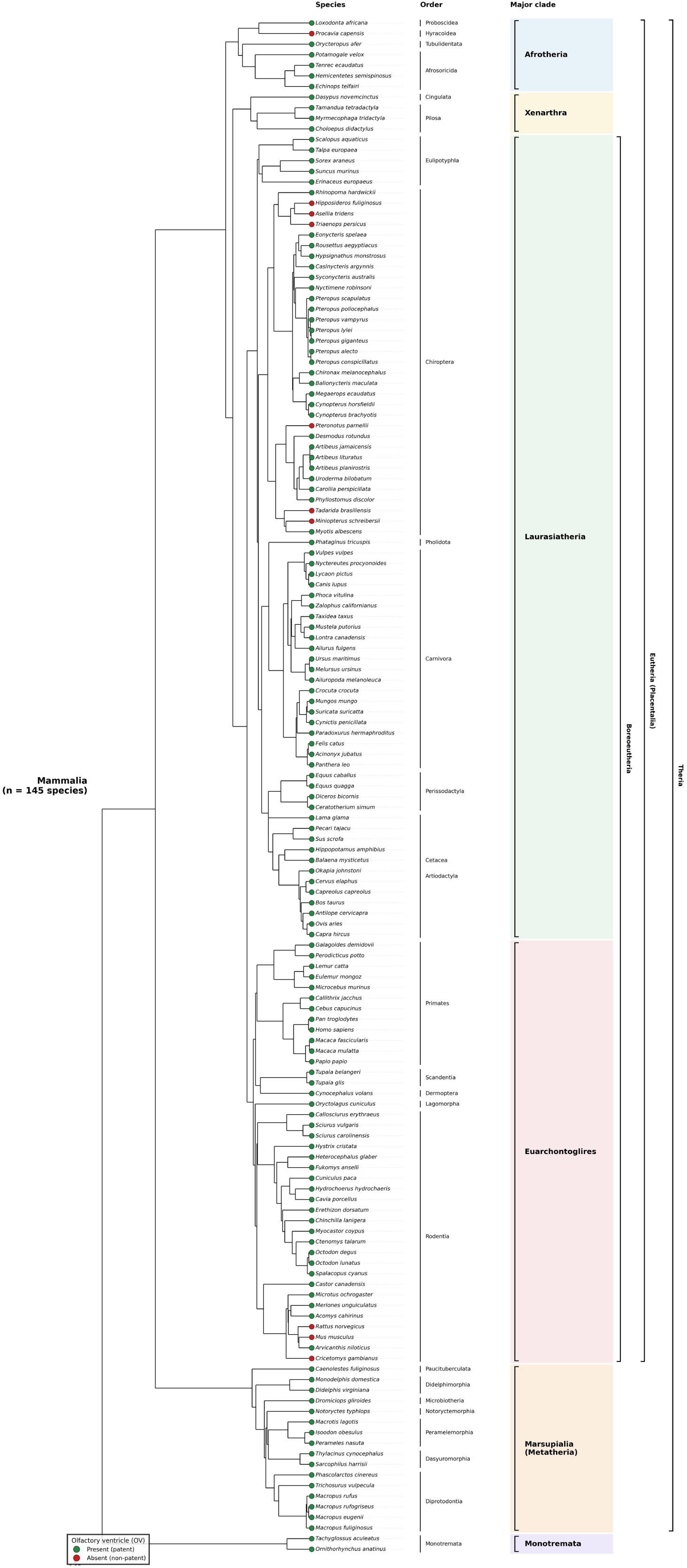
Phylogenetic distribution of the OV across Mammalia. Phylogenetic distribution of OV status across 145 mammalian species. Species were classified according to the presence of a patent OV in adulthood (green circles) or the absence of a patent OV (red circles). Species are organized according to the mammalian phylogeny of Upham et al. (2019), with taxonomic orders and major mammalian clades indicated on the right. Higher-level brackets indicate Boreoeutheria, Eutheria (Placentalia), and Theria. OV persistence is broadly distributed across Mammalia, whereas absence of a patent OV is restricted to a limited number of lineages.

Within Hyracoidea, the single sampled species, *Procavia capensis*, lacked a patent OV. Among Rodentia, an open OV was observed in 24 of 27 observations, whereas absence was recorded in *Cricetomys gambianus*, C57BL/6 *Mus musculus* and Wistar *Rattus norvegicus*. Importantly, this pattern was not uniform within the latter two species, as an open OV was observed in other *M. musculus* and *R. norvegicus* populations included in the survey, revealing intraspecific variation in adult OV persistence.

Chiroptera showed the greatest taxonomic diversity in OV status. An open OV was observed in 28 of 34 sampled bats, whereas six lacked a patent OV: *Hipposideros fuliginosus* and *Asellia tridens* (Hipposideridae), *Triaenops persicus* (Rhinonycteridae), *Miniopterus schreibersii* (Miniopteridae), *Tadarida brasiliensis* (Molossidae), and *Pteronotus parnellii* (Mormoopidae). Conversely, OV presence was consistently observed in several extensively sampled bat groups, including all sampled Pteropodidae and Phyllostomidae.

Together, these observations reveal that the OV is broadly distributed across mammalian diversity, whereas its absence is comparatively rare and concentrated within a restricted subset of taxonomic lineages.

### 3.2. Phylogenetic reconstruction supports an ancestral OV followed by secondary losses

To investigate the evolutionary history underlying this taxonomically restricted pattern of OV absence, we reconstructed ancestral OV states across 100 alternative mammalian phylogenies. Following taxonomic reconciliation and aggregation of observations corresponding to the same phylogenetic species, the comparative dataset comprised 145 unique phylogenetic tips (Fig. 1). In the primary analysis, *Mus musculus* and *Rattus norvegicus* were conservatively coded as OV-absent because OV absence was documented in at least one population of each species; the alternative coding was explicitly evaluated in sensitivity analyses.

The all-rates-different (ARD) model, allowing gains and losses of the OV to occur at different rates, provided a better fit to the data than the equal-rates (ER) model. The median difference in AIC between the ER and ARD models (ΔAICER-ARD) was 7.45, with the ARD model favored by ΔAIC > 2 across all 100 phylogenies. Under the ARD model, ancestral-state reconstruction consistently supported the presence of an OV in the mammalian ancestor. The median probability of OV presence at the root was 0.948, with values ranging from 0.941 to 0.954 across the central 95% of the sampled phylogenies.

This inference was robust to the intraspecific variability observed in *M. musculus* and *R. norvegicus*. When both species were alternatively coded as OV-present, the median probability of OV presence at the mammalian root remained 0.953 (central 95% of phylogenies: 0.944-0.969). Under this alternative coding, the ARD model remained favored over the ER model, with a median ΔAICER-ARD of 4.15 and ΔAIC > 2 in 88 of the 100 phylogenies. Thus, uncertainty in the coding of these two species had little effect on inference of the ancestral mammalian state.

Given the strong support for OV presence in the mammalian ancestor, maximum-parsimony reconstruction was subsequently used to identify the minimum number of transitions required to account for the observed distribution. With *M. musculus* and *R. norvegicus* coded as OV-absent, the observed pattern could be explained by a minimum of eight independent losses, with no secondary gain required, across all 100 phylogenies. Four of these losses occurred within Chiroptera (see below).

Together, the ancestral-state and parsimony reconstructions support an evolutionary scenario in which the OV represents an ancestral and broadly conserved feature of the mammalian brain that has undergone a limited number of secondary losses during mammalian diversification.

### 3.3. OV distribution exhibits a strong phylogenetic signal

The restricted distribution of OV absence suggested that evolutionary losses might be phylogenetically clustered rather than randomly distributed across mammals. We therefore quantified phylogenetic signal using the D statistic for binary traits. With *Mus musculus* and *Rattus norvegicus* coded as OV-absent, median D across the 100 phylogenies was -0.071, with values ranging from -0.125 to 0.010 across the central 95% of trees. In all 100 phylogenies, the observed distribution differed significantly from that expected under a random association of OV states with the phylogeny (p < 0.05). By contrast, none of the phylogenies showed a significant departure from the phylogenetic structure expected under a Brownian threshold model (median p = 0.595).

The same conclusion was obtained when *M. musculus* and *R. norvegicus* were alternatively coded as OV-present. Under this scenario, phylogenetic clustering was even stronger (median D = -0.244; central 95% of trees: -0.367 to -0.119), with significant departure from phylogenetic randomness across all 100 trees but no significant departure from the Brownian expectation.

These analyses show that OV absence is not randomly distributed among mammalian species but instead exhibits a strong phylogenetic structure, consistent with losses concentrated within particular evolutionary lineages.

### 3.4. Repeated independent losses of the OV occurred within Chiroptera

Chiroptera represented the mammalian order with the greatest diversity in OV status and accounted for six of the ten OV-absent observations in the comparative dataset. Mapping OV status onto the mammalian phylogenies revealed that these absences were not randomly interspersed among OV-present species but were concentrated within a limited number of bat lineages.

Maximum-parsimony reconstruction supported four independent losses within Chiroptera across all 100 phylogenies (Fig. 2). A single loss was sufficient to explain OV absence in the sampled Hipposideridae-Rhinonycteridae lineage, comprising *Hipposideros fuliginosus*, *Asellia tridens* and *Triaenops persicus*. Three additional losses were independently reconstructed along the lineages leading to *Pteronotus parnellii* (Mormoopidae), *Tadarida brasiliensis* (Molossidae) and *Miniopterus schreibersii* (Miniopteridae).

**Figure 2.**
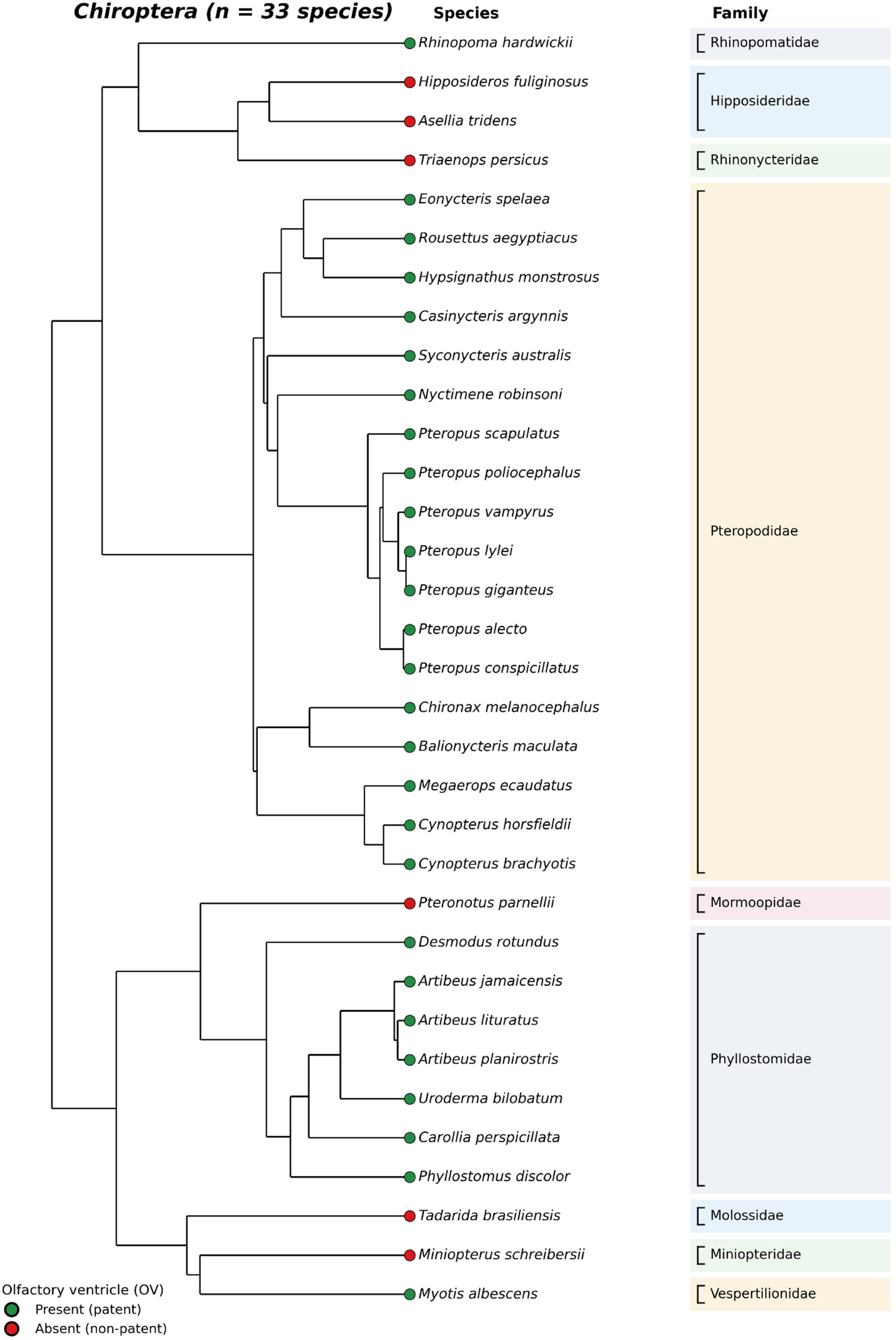
Phylogenetic distribution of the OV within Chiroptera. Phylogenetic distribution of OV status across 33 chiropteran species. Green circles indicate the presence of a patent OV and red circles the absence of a patent OV. Family affiliations are indicated on the right. OV absence is distributed across several phylogenetically distinct chiropteran lineages.

**Figure 3.**
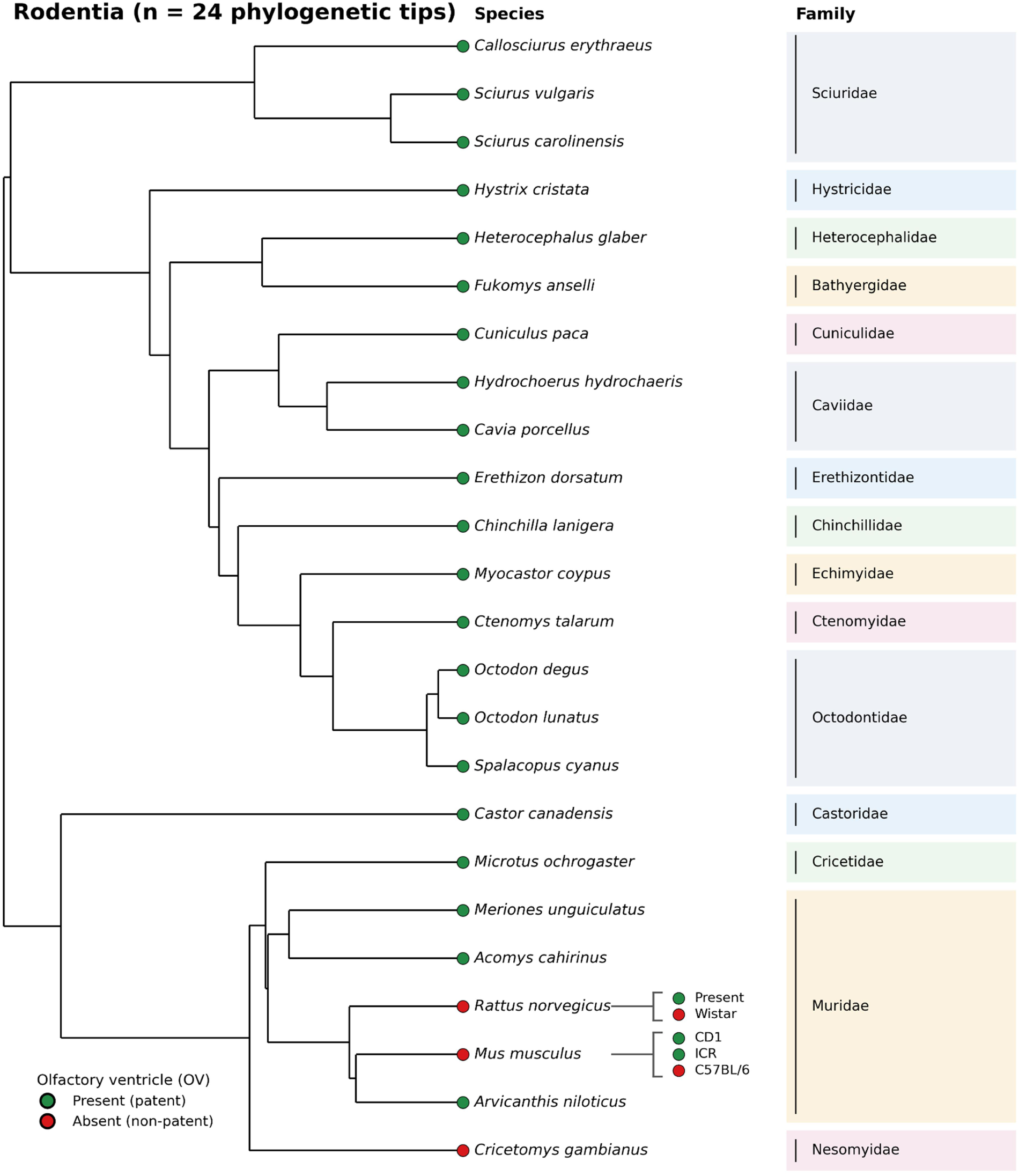
Phylogenetic distribution of the OV within Rodentia and intraspecific variation in *Mus musculus* and *Rattus norvegicus*. Phylogenetic distribution of OV status across 24 rodent phylogenetic tips. Green circles indicate the presence of a patent OV and red circles the absence of a patent OV. Family affiliations are indicated on the right. For *Rattus norvegicus* and *Mus musculus*, for which contrasting observations were available, individual observations are shown adjacent to the corresponding species. A patent OV was observed in *R. norvegicus*, whereas it was absent in Wistar rats. In *M. musculus*, the OV was patent in CD1 and ICR mice but non-patent in C57BL/6 mice. The circle at each phylogenetic tip indicates the species-level state used in the primary phylogenetic analysis.

In contrast, the OV was retained across several other chiropteran lineages, including all sampled Pteropodidae and Phyllostomidae. The coexistence of deeply conserved OV presence and repeated lineage-specific losses within a single mammalian order highlights Chiroptera as a particularly informative group for understanding the evolutionary diversification of olfactory ventricular anatomy.

### 3.5. OV persistence varies at the intraspecific level within Muridae

Muridae revealed a distinct pattern of OV variation compared with the lineage-specific losses observed in Chiroptera. Eight anatomical observations representing five murid species were included in the comparative dataset. An open OV was consistently observed in *Arvicanthis niloticus*, *Acomys cahirinus* and *Meriones unguiculatus*. By contrast, both *Mus musculus* and *Rattus norvegicus* displayed intraspecific variation in adult OV persistence.

Among *M. musculus*, an open OV was observed in CD1 and ICR mice, whereas no patent OV was detected in C57BL/6 mice. Similarly, an open OV was documented in *R. norvegicus*, whereas absence was reported in Wistar rats. Thus, OV status was not uniform at the species level in either *Mus* or *Rattus*. Unlike the deeper lineage-associated pattern observed in Chiroptera, OV variation within Muridae included variation at the intraspecific level. Instead, the coexistence of OV-present and OV-absent populations within the same species indicates that closure of the adult OV can vary at an evolutionarily recent, intraspecific scale.

### 3.6. Cellular organization of the persistent olfactory ventricle across mammalian species

Histological examination revealed marked interspecific differences in the morphology of the persistent OV. In adult sheep, the OV formed a large and clearly patent ventricular cavity bordered by a well-defined ventricular wall (Fig. 4A-C). In Peromyscus maniculatus, the OV was retained in adulthood but displayed a markedly narrower, collapsed configuration, with closely apposed ventricular walls and a restricted lumen (Fig. 4D-F). In adult CD1 mice, the OV remained readily identifiable and was surrounded by a particularly thick and densely cellular periventricular region (Fig. 4G-I). Thus, despite its persistence in all three species, the adult OV exhibited substantial variation in lumen configuration and surrounding wall morphology.

**Figure 4.**
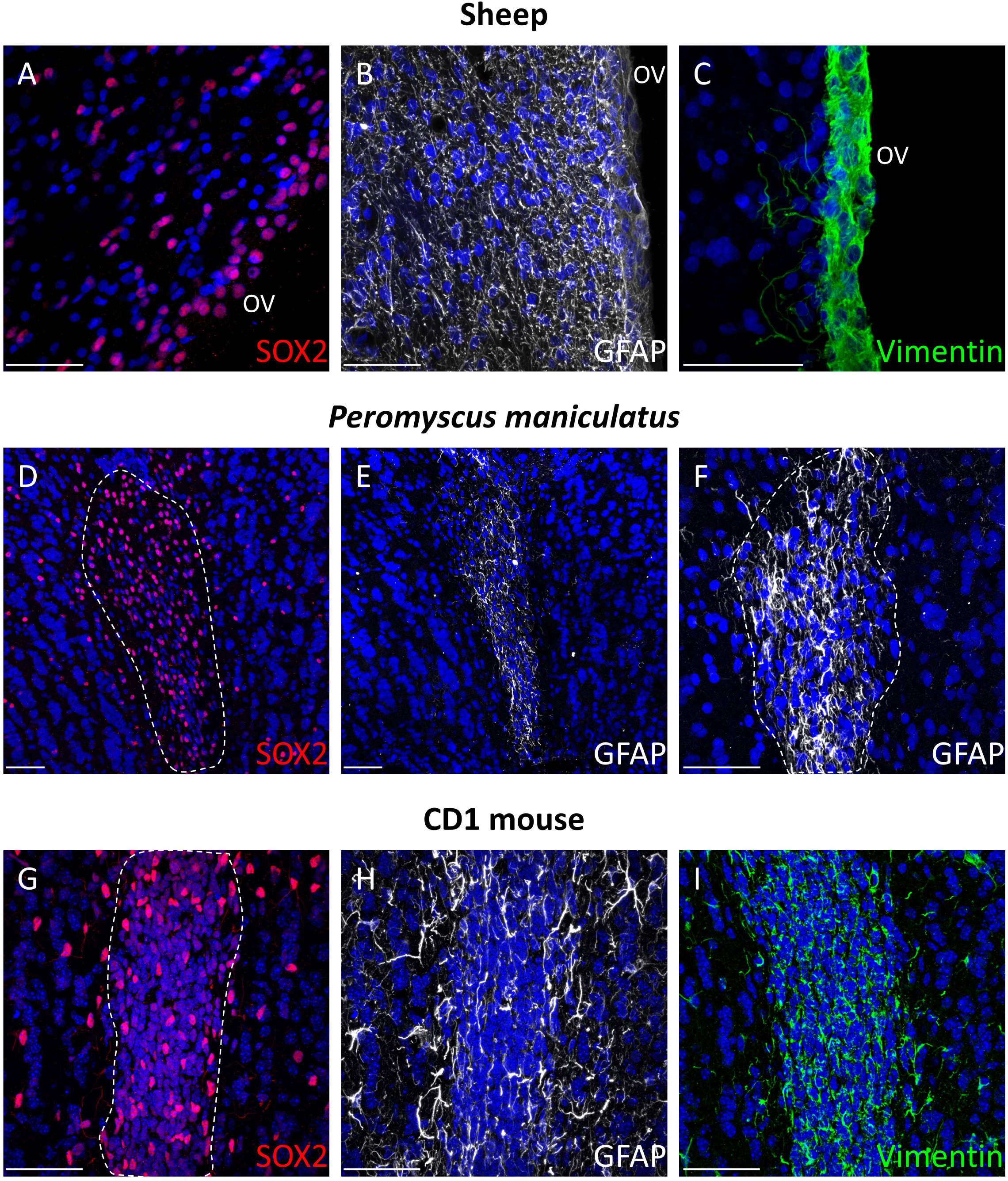
Progenitor- and glial-associated features of the adult olfactory ventricular region across mammalian species. Representative confocal images of the adult OV and surrounding periventricular region in sheep (**A-C**), *Peromyscus maniculatus* (**D-F**), and CD1 mice (**G-I**). SOX2 immunoreactivity identifies abundant cells associated with the OV region in sheep (**A**), *P. maniculatus* (**D**), and CD1 mice (**G**). GFAP immunoreactivity reveals distinct species-specific distributions associated with the OV region in sheep (B), *P. maniculatus* (E-F), and CD1 mice (H) Vimentin immunoreactivity further delineates the OV-associated region in sheep (**C**) and CD1 mice (**I**). Nuclei are counterstained with DAPI. OV, olfactory ventricle. Scale bar, 50 µm.

To determine whether this anatomical diversity was accompanied by conserved cellular features, we first examined the distribution of progenitor- and glial-associated markers (Fig. 4). SOX2-positive cells were consistently associated with the OV region in sheep, *P. maniculatus* and CD1 mice, although their organization differed among species. In sheep, SOX2-positive nuclei were concentrated along the ventricular wall (Fig. 4A), whereas in *P. maniculatus* they delineated the closely apposed borders of the collapsed OV (Fig. 4D). In CD1 mice, numerous SOX2-positive cells occupied the thick periventricular region surrounding the OV (Fig. 4G). GFAP immunoreactivity was also associated with the OV region in all three species, although its spatial distribution differed markedly. In sheep, GFAP immunoreactivity was relatively diffuse throughout the tissue surrounding the OV. In P. maniculatus, by contrast, GFAP immunoreactivity was almost exclusively concentrated along the OV border. In CD1 mice, GFAP immunoreactivity was predominantly concentrated within the OV-associated periventricular region, with additional staining extending into the surrounding olfactory bulb parenchyma. Vimentin further delineated the OV-associated region in sheep and CD1 mice (Fig. 4C, I). Together, these observations reveal a progenitor- and glial-marker-rich cellular organization associated with the persistent adult OV despite substantial interspecific differences in ventricular morphology.

### 3.7. Immature neuronal and proliferative cells are associated with the adult OV

We next examined markers of immature neurons and cell proliferation (Fig. 5). DCX-positive cells were detected in association with the OV in all three species examined. In sheep, rounded DCX-positive cells were present within and immediately adjacent to the ventricular wall (Fig. 5A), whereas the olfactory bulb parenchyma displayed a markedly different pattern, characterized by extensive fibrillary DCX immunoreactivity and numerous DCX-positive processes (Fig. 5B). In *P. maniculatus*, small rounded DCX-positive cells lacking obvious processes were concentrated at the OV border (Fig. 5D), whereas DCX-positive cells located farther within the olfactory bulb parenchyma displayed more elongated or ovoid morphologies and cellular processes (Fig. 5E). In CD1 mice, DCX immunoreactivity was abundant within the OV-associated periventricular region (Fig. 5F), while DCX-positive cells were also observed within the olfactory bulb parenchyma (Fig. 5G). These observations identify immature neuronal populations in close spatial association with the persistent OV, although their site of origin cannot be determined from DCX immunoreactivity alone.

**Figure 5.**
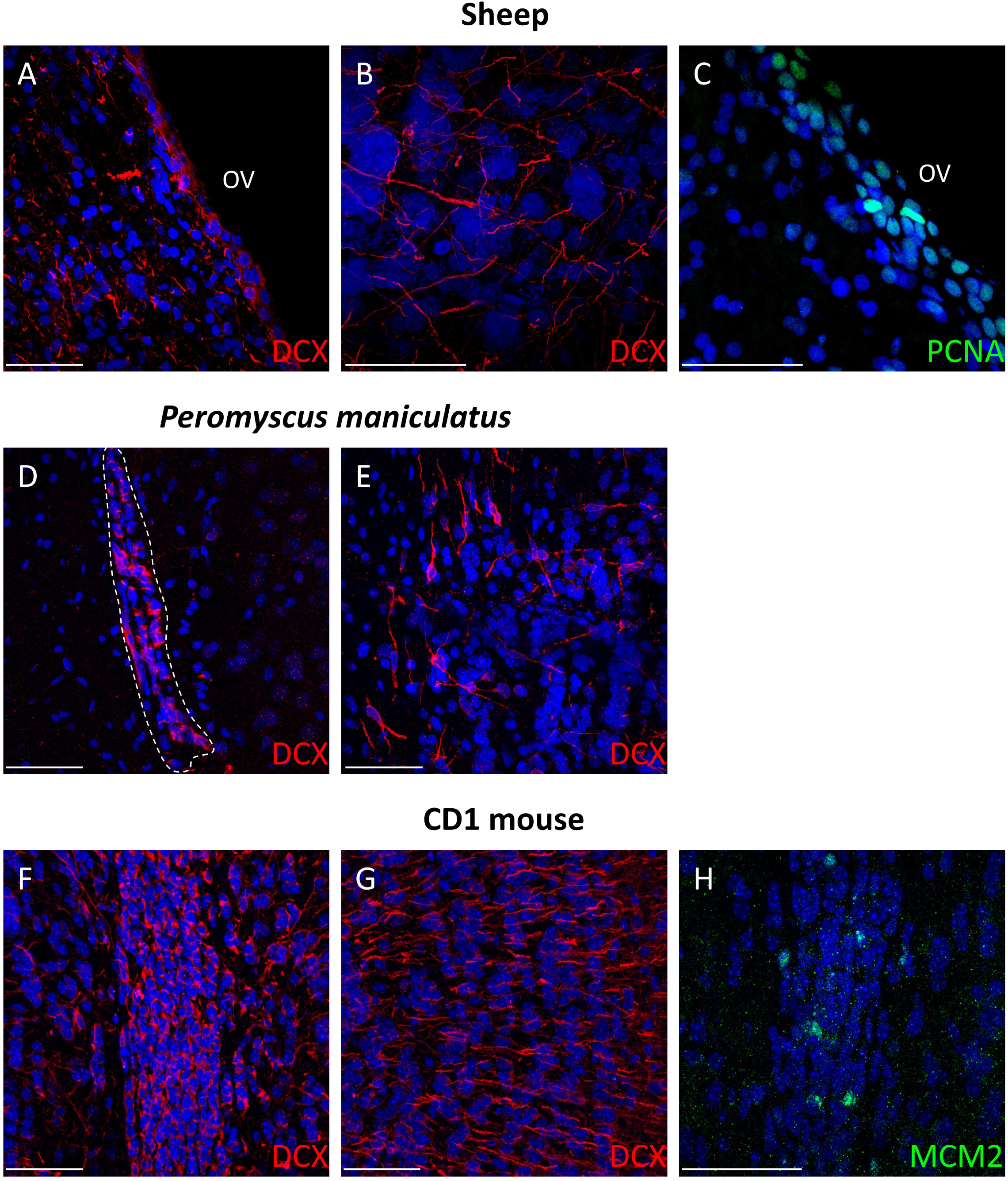
Proliferative and immature neuronal populations associated with the adult OV. Representative confocal images of proliferative and DCX-positive cells in the OV-associated region of sheep (**A-C**), *Peromyscus maniculatus* (**D-E**), and CD1 mice (**F-H**). In sheep, DCX immunoreactivity identifies immature neuronal cells within the OV-associated region and surrounding olfactory bulb parenchyma (**A-B**), while PCNA-positive cells are present along the ventricular wall (**C**). In *P. maniculatus*, DCX-positive cells are observed in close association with the collapsed OV (**D**) and in the surrounding tissue (**E**), displaying different cellular morphologies. In CD1 mice, DCX-positive cells are abundant within the OV-associated periventricular region (**F**) and olfactory bulb parenchyma (**G**), while MCM2-positive cells provide evidence of proliferative activity within the OV (**H**). Nuclei are counterstained with DAPI. OV, olfactory ventricle. Scale bar, 50 µm.

In sheep, PCNA-positive nuclei were detected along the OV wall (Fig. 5C), indicating the presence of proliferative cells within the ventricular region. In CD1 mice, MCM2-positive nuclei were similarly observed within the OV-associated periventricular region, providing independent evidence that proliferative cells are retained in association with the persistent OV (Fig. 5H).

Together, the presence of proliferating cells in close association with the OV wall and of DCX-positive cells displaying immature neuronal morphologies supports the presence of cellular features associated with adult neurogenesis around the persistent OV. The spatial change in DCX-positive cell morphology was consistent with progressive neuronal maturation with increasing distance from the ventricular border.

### 3.8. The persistent OV retains an organized apical and ciliary-associated interface

We next examined proteins associated with ventricular polarity and ciliary organization (Figs. 6-8). Despite marked differences in OV morphology, all three species displayed a molecularly organized ventricular interface. In sheep, β-catenin and CD133 immunoreactivity delineated the ventricular border, consistent with the maintenance of a specialized apical domain (Fig. 6A-B). Acetylated α-tubulin was strongly enriched at the ventricular border and was additionally detected in fibrillary structures within the surrounding parenchyma (Fig. 6C). ARL13B-positive structures were detected at the ventricular interface (Fig. 6D), while AC3 displayed a punctate distribution along the OV wall (Fig. 6E-F).

**Figure 6.**
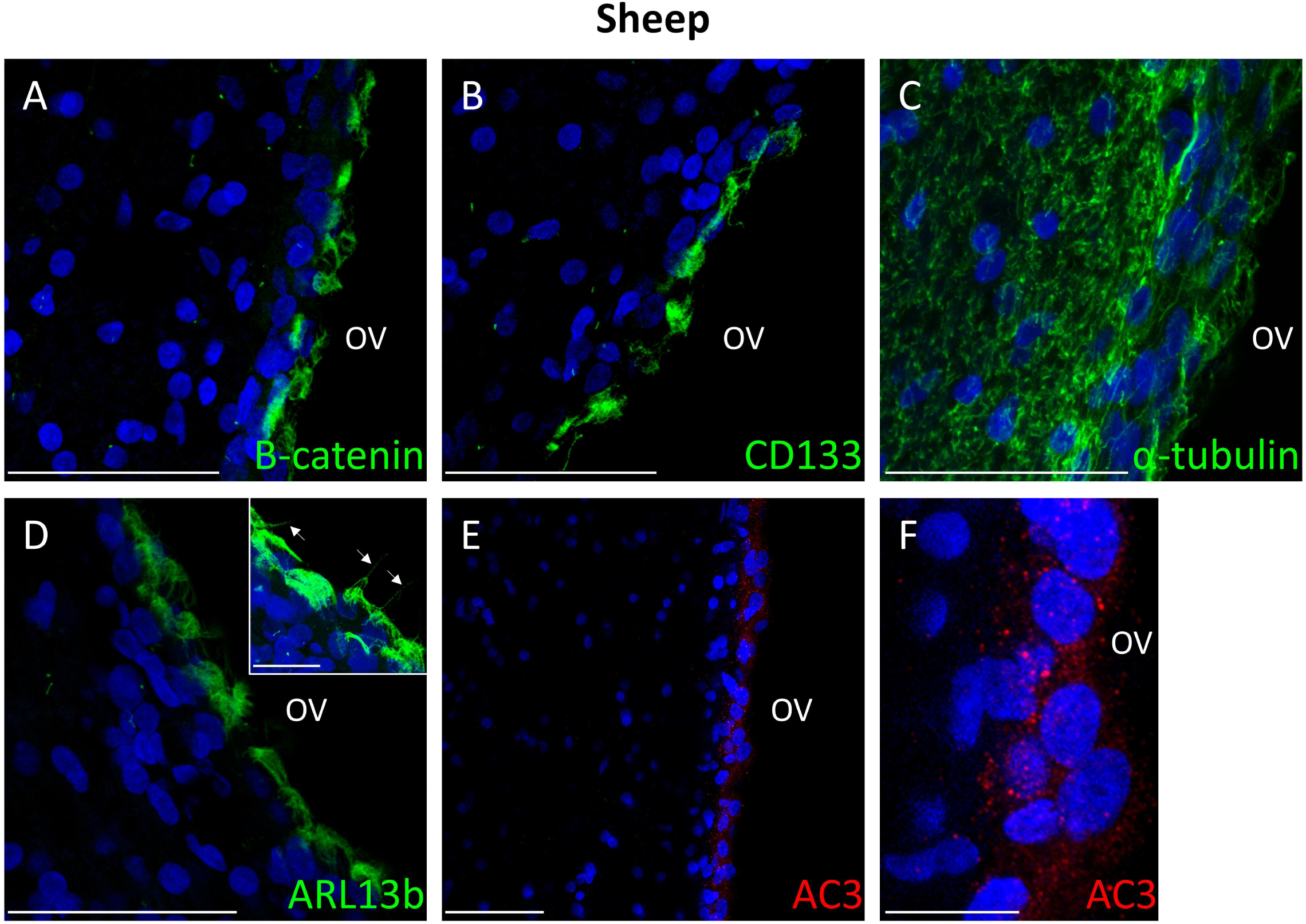
Apical and ciliary-associated organization of the adult olfactory ventricular wall in sheep. Representative confocal images showing the distribution of markers associated with ventricular epithelial organization and ciliary structures along the adult sheep OV. β-catenin immunoreactivity delineates the ventricular border (**A**), while CD133 (**B**) and ARL13B (**D**) immunoreactivity is associated with the apical ventricular region. Acetylated α-tubulin staining highlights structures along the ventricular surface and fibrillary elements extending into the adjacent tissue (**C**). AC3 immunoreactivity is detected as discrete punctate structures associated with the ventricular wall (**E-F**). Nuclei are counterstained with DAPI. OV, olfactory ventricle. Scale bar, 50 µm, 20 µm (D) and 10 µm (F).

A comparable organization was observed in *P. maniculatus* (Fig. 7). β-catenin immunoreactivity delineated the closely apposed OV borders (Fig. 7A-B), while CD133 was detected in association with the OV (Fig. 7C). ARL13B-positive structures and punctate AC3 immunoreactivity were also associated with the ventricular region (Fig. 7D-F). Despite the markedly collapsed morphology of the ventricle in this species, these markers revealed the persistence of a molecularly differentiated ventricular interface.

**Figure 7.**
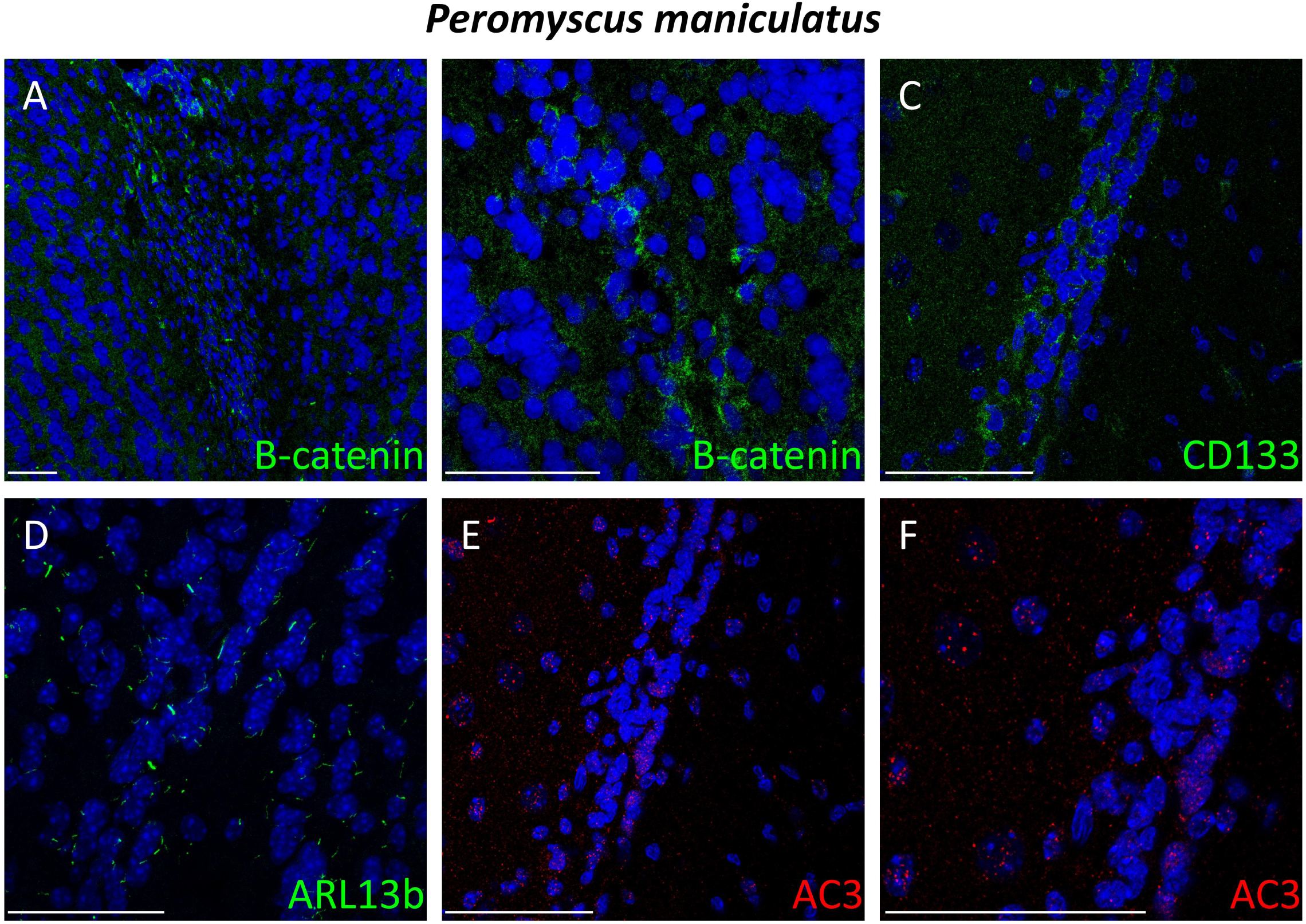
Apical and ciliary-associated features of the adult olfactory ventricular region in *Peromyscus maniculatus*. Representative confocal images of the OV-associated region in adult *P. maniculatus*. β-catenin immunoreactivity is detected within the OV-associated region (**A-B**). CD133 (**C**) and ARL13B (**D**) immunoreactivity further reveal molecular features associated with ventricular/apical organization. AC3-positive structures are detected within the same region (**E-F**), including discrete punctate profiles. Nuclei are counterstained with DAPI. Scale bar, 50 µm.

CD1 mice similarly retained several features associated with apical and ciliary organization (Fig. 8). β-catenin immunoreactivity was associated with the OV region (Fig. 8A), acetylated α-tubulin was enriched at the ventricular interface (Fig. 8B-C), and ARL13B-positive structures were detected in the ventricular region (Fig. 8D). AC3 immunoreactivity displayed a punctate pattern associated with the OV (Fig. 8E-F). Together, the enrichment of apical-associated proteins at the ventricular border and the presence of multiple ciliary-associated markers reveal a specialized apical organization of the persistent OV wall across the mammalian species examined.

**Figure 8.**
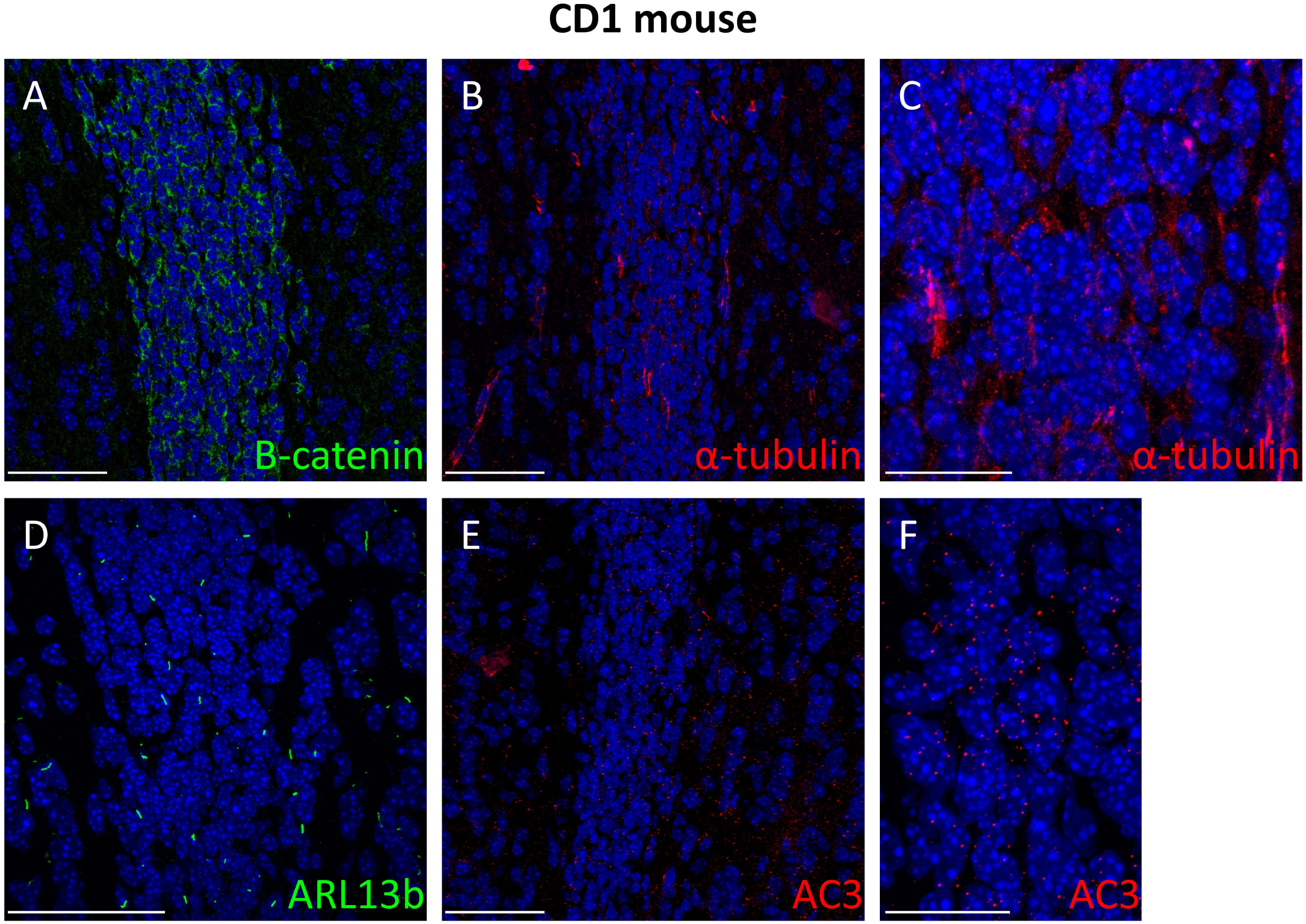
Apical and ciliary-associated features of the adult olfactory ventricular region in CD1 mice. Representative confocal images showing markers associated with apical and ciliary organization in the OV-associated region of adult CD1 mice. β-catenin immunoreactivity is detected throughout the OV-associated region (**A**). Acetylated α-tubulin staining reveals prominent fibrillary and punctate structures (**B-C**), while ARL13B immunoreactivity identifies discrete ciliary-associated profiles (**D**). AC3 immunoreactivity is also detected within the OV- associated region (**E-F**), including numerous punctate structures. Nuclei are counterstained with DAPI. Scale bars, 50 µm and 20 µm (F).

### 3.9. Multiplex staining identifies proliferating progenitor-associated cells at the sheep OV wall

Finally, multiplex immunofluorescence was used in sheep to determine whether progenitor, glial and proliferative markers were expressed within the same ventricular cell populations (Fig. 9). Triple labelling for SOX2, GFAP and vimentin revealed cells coexpressing the three markers along the ventricular border (Fig. 9A). These triple-labelled cells were concentrated within the OV wall, identifying a population with a progenitor/glial phenotype lining the adult ventricular compartment. In a second combination, SOX2-positive nuclei expressing PCNA were detected within the vimentin-positive ventricular region (Fig. 9B). These observations demonstrate that proliferative cells expressing progenitor-associated markers are located within the specialized cellular compartment bordering the adult sheep OV.

**Figure 9.**
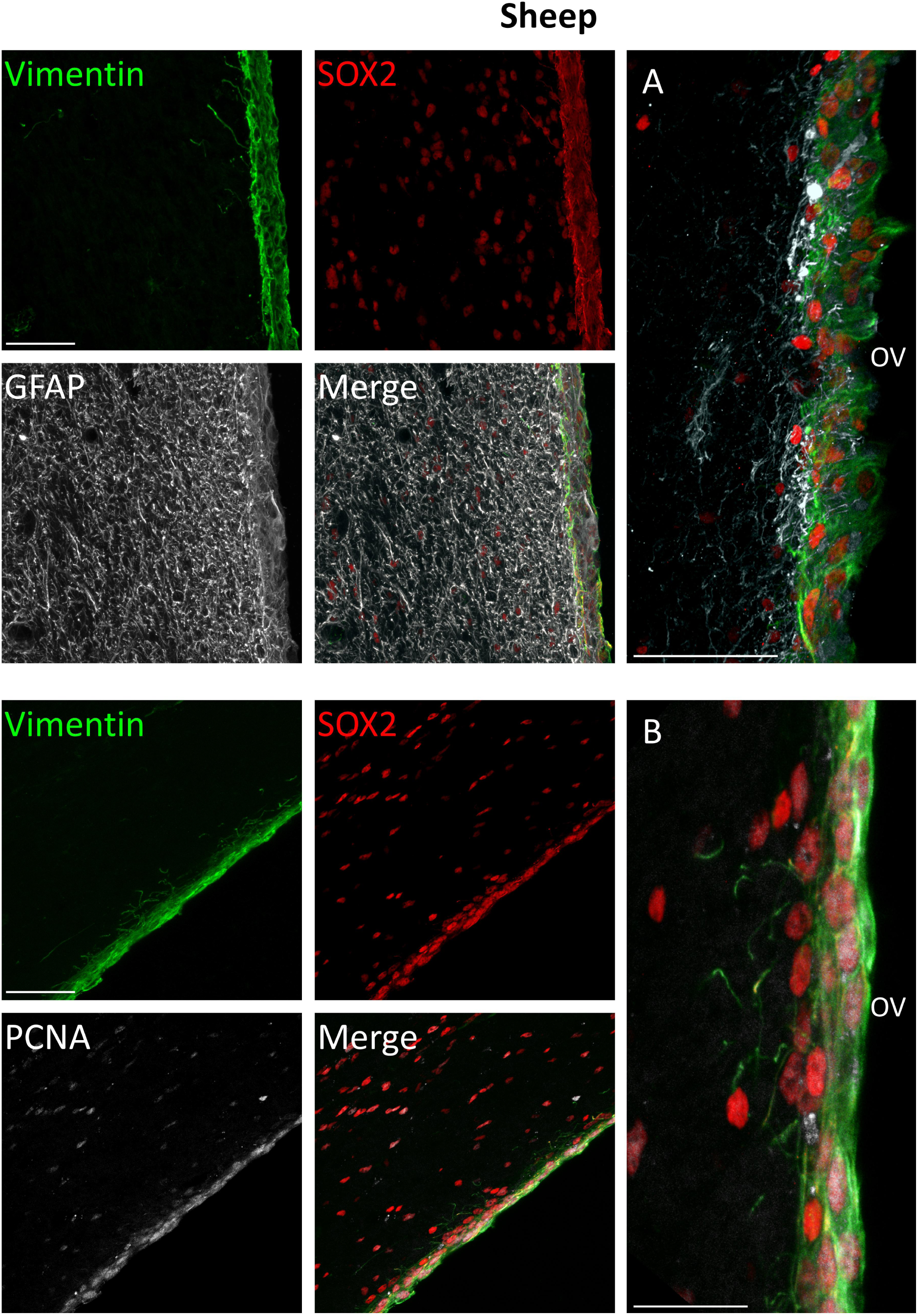
Multiplex characterization of the cellular organization of the adult sheep olfactory ventricular wall. Multiplex immunofluorescence analysis of the adult sheep OV wall. **(A)** Combined detection of vimentin, SOX2, and GFAP reveals a densely organized ventricular region in which SOX2-positive nuclei are embedded within a vimentin- and GFAP-associated cellular network. Individual channels and the merged image are shown. **(B)** Combined detection of vimentin, SOX2, and PCNA identifies proliferative PCNA-positive cells within the SOX2- and vimentin-associated ventricular region. Individual channels and the merged image are shown. Higher-magnification views illustrate the spatial association of these markers at the ventricular border. OV, olfactory ventricle. Scale bars, 50 µm.

## 4. Discussion

Our comparative analysis substantially revises the prevailing view of the OV as a transient structure that disappears during postnatal development. Across the mammalian diversity examined here, a patent adult OV was observed in the great majority of anatomical observations and across all major mammalian lineages, including monotremes, marsupials and placental mammals. Conversely, absence of a patent adult OV was restricted to a small number of taxa within Hyracoidea, Rodentia and Chiroptera. Persistence of the OV into adulthood therefore appears not to represent an exceptional anatomical condition restricted to a few mammalian groups, but rather a widespread feature of mammalian OB organization. This finding is particularly relevant because current descriptions of OV development have been strongly influenced by laboratory rodents, and especially by Mus musculus strains in which the olfactory ventricular cavity closes during early postnatal development.

The broad taxonomic distribution of the OV is consistent with an ancient evolutionary origin. Ancestral-state reconstruction across 100 alternative mammalian phylogenies consistently supported OV presence at the mammalian root, while models allowing asymmetric rates of OV gain and loss were better supported than equal-rate models. Importantly, this inference was robust to alternative coding of Mus musculus and Rattus norvegicus, the two species for which both OV-present and OV-absent observations were identified. These analyses therefore support a scenario in which a patent OV was present in the mammalian ancestor, with its absence in extant species predominantly reflecting secondary loss rather than repeated independent acquisition.

Chiroptera provide a particularly informative case for investigating the evolutionary factors associated with OV loss. All six OV-absent species identified in our dataset - *Hipposideros fuliginosus, Asellia tridens, Triaenops persicus, Pteronotus parnellii, Tadarida brasiliensis and Miniopterus schreibersii* - are predominantly insectivorous and rely extensively on echolocation for prey detection, whereas the OV was retained in several sampled lineages in which olfaction plays a prominent role in foraging, notably Pteropodidae and Phyllostomidae. This pattern is consistent with comparative evidence showing that bat sensory systems have diversified in association with feeding ecology, with greater investment in olfactory structures in plant-feeding bats and increased specialization of auditory systems in aerial insectivores (Bhatnagar & Kallen, 1974; Hutcheon et al., 2002). More recent comparative analyses have further suggested an evolutionary trade-off between olfactory and echolocation-related specializations (López-Aguirre et al., 2025). This raises the possibility that repeated OV loss in Chiroptera is associated with shifts in sensory ecology, with increased reliance on echolocation potentially relaxing selective constraints on the maintenance of olfactory ventricular architecture. However, broader sampling and phylogenetically controlled analyses will be required to test this relationship.

OV loss does not appear to imply loss of adult olfactory neurogenesis, but may instead be associated with a different spatial organization of neuronal production. Chawana et al. (2016) examined three of the OV-absent species identified here - *H. fuliginosus, T. persicus and A. tridens* - and reported proliferating Ki-67-positive cells and DCX-positive immature neuronal populations within the SVZ, together with a prominent RMS extending toward the olfactory bulb and abundant DCX-positive cells within the bulb. This organization is consistent with a prominent SVZ-RMS-OB neurogenic pathway in these OV-negative species. Thus, OV-negative mammals may rely predominantly on progenitors generated at a distance in the SVZ and recruited to the olfactory bulb through the RMS, whereas persistence of the OV could potentially provide an additional local progenitor compartment directly within the bulb. Under this model, evolutionary variation in OV persistence would influence the spatial organization rather than simply the presence or absence of adult olfactory neurogenesis. This interpretation remains hypothetical, however, as neither the origin of the DCX-positive cells described by Chawana et al. nor local neuronal production from OV-associated progenitors has been directly demonstrated. Comparative lineage-tracing experiments will therefore be required to test the relative contribution of SVZ-derived and local progenitors in OV-negative and OV-positive mammals.

Conversely, variation within Muridae extends to the intraspecific level, demonstrating that the developmental fate of the OV can differ even among closely related populations. These two patterns suggest that adult OV persistence is not constrained by a single evolutionary trajectory but may be modified independently through changes in the developmental processes governing maintenance or closure of the anterior ventricular system.

In *M. musculus*, the only population in our dataset lacking a patent adult OV was C57BL/6, a highly inbred laboratory strain, whereas the OV persisted in the outbred CD1 and ICR stocks. This pattern raises the possibility that OV persistence may be influenced by genetic background. Substantial strain-dependent variation in mouse brain anatomy has been demonstrated by MRI, including differences between the inbred C57BL/6 and 129S6 strains and the outbred CD1 stock (Scholz et al., 2016). Genetic background also influences several aspects of adult neurogenesis (Kempermann et al., 1997), and, particularly relevant to the olfactory system, marked differences in neuroblast migration along the SVZ-RMS and olfactory sensitivity have been reported among C57BL/6, BALB/c and 129/S1 mice (Lee et al., 2003). Together, these studies demonstrate that genetic background can substantially influence structural and plasticity-related features of the mouse forebrain and olfactory system. Whether OV closure in C57BL/6 represents another strain-specific developmental phenotype remains unknown.

Importantly, our observations cannot distinguish an effect of inbreeding per se from the effects of particular genetic variants fixed in the C57BL/6 lineage. Examination of additional inbred strains, outbred stocks and wild-derived M. musculus populations will therefore be necessary to determine whether adult OV persistence is associated with breeding history and genetic diversity or instead reflects a developmental trajectory specific to particular laboratory strains. More generally, the developmental mechanisms responsible for evolutionary or intraspecific variation in adult OV persistence remain unknown. Absence of a patent adult OV need not imply absence of an olfactory ventricular structure throughout development. Instead, it may result from variation in the timing or completeness of postnatal ventricular closure. In the laboratory mouse, the embryonic olfactory ventricular zone constitutes the rostral extension of the LGE, and progressive collapse and fusion of the OV walls during the early postnatal period accompanies formation of the RMS (Jacquet et al., 2011). The canonical adult SVZ-RMS-OB organization of C57BL/6 mice can therefore be viewed as the outcome of substantial postnatal remodeling of an initially ventricular neurogenic system. Whether comparable remodeling occurs in other OV-negative mammals, and whether evolutionary retention of the adult OV results from incomplete execution or active maintenance of this developmental program, remain unknown. Comparative developmental analyses across OV-present and OV-absent taxa will be essential to distinguish these possibilities.

The widespread evolutionary retention of the OV also raises a more fundamental question: does persistence of this structure have biological significance beyond representing a remnant of embryonic ventricular anatomy? Anatomical conservation alone cannot establish function. However, the preservation of a structured cellular environment in phylogenetically and anatomically distinct species argues against viewing the adult OV simply as an empty residual cavity. Our histological observations instead reveal several features that are characteristic of ventricular regions associated with neural progenitor populations.

SOX2-positive cells were consistently associated with the OV in sheep, *P. maniculatus* and CD1 mice, together with GFAP immunoreactivity, although the organization of the latter differed markedly among species. GFAP was relatively diffuse around the sheep OV, became strongly restricted to the ventricular border in P. maniculatus, and was predominantly concentrated in the thick periventricular region of CD1 mice while extending to a lesser extent into the surrounding parenchyma. Vimentin additionally delineated the OV-associated region in both sheep and CD1 mice. Thus, the conserved feature is not a rigidly identical ventricular wall but rather the association of the persistent OV with a progenitor- and glial-marker-rich cellular compartment whose spatial organization varies substantially among species (Doetsch et al., 1999; Mirzadeh et al., 2008). Adult neurogenic niches themselves exhibit considerable anatomical and cellular heterogeneity across mammals while retaining common features related to progenitor maintenance and ventricular organization (Bonfanti et al., 2024; Paredes et al., 2016).

Proliferative cells were also associated with the persistent OV. In sheep, multiplex labelling identified SOX2/PCNA-positive cells within the vimentin-rich ventricular compartment, directly linking proliferative activity to cells expressing a progenitor-associated marker at the OV border. In CD1 mice, MCM2-positive cells were likewise detected within the OV-associated periventricular region. The use of distinct proliferation markers in two species precludes a direct quantitative comparison, but their convergence indicates that proliferative cells are not restricted to a single species with an unusually large OV. Together with the SOX2-, GFAP- and vimentin-associated organization, a combination characteristic of neural stem/progenitor populations in adult ventricular niches (Doetsch et al., 1999; van Strien et al., 2014), these observations support the presence of proliferating neural stem/progenitor-like populations associated with the persistent OV. Their proliferative capacity and lineage potential, however, remain to be established experimentally.

DCX-positive immature neuronal populations were also consistently associated with the persistent OV, although their organization differed among species. Importantly, their organization was not identical among species. In sheep, rounded DCX-positive cells were observed at and immediately adjacent to the ventricular wall, whereas the surrounding OB parenchyma displayed extensive fibrillary DCX immunoreactivity. In P. maniculatus, the contrast was more suggestive of a morphological progression: small rounded DCX-positive cells lacking obvious processes were concentrated at the OV border, whereas cells farther within the OB parenchyma displayed more elongated or ovoid morphologies and cellular processes. CD1 mice also contained abundant DCX-positive cells in the OV-associated periventricular region, together with DCX immunoreactivity within the OB parenchyma. The recurrence of immature neuronal markers in close spatial association with the OV across all three species is notable, particularly when considered together with the progenitor-associated and proliferative populations described above. In P. maniculatus, the spatial change in DCX-positive cell morphology is compatible with progressive neuronal maturation away from the ventricular border. Nevertheless, DCX expression identifies neuronal immaturity rather than neuronal birth per se, and DCX-positive populations persisting into adulthood may include prenatally generated neurons, as demonstrated in sheep (Piumatti et al., 2018). Neither DCX expression nor cellular morphology can therefore establish where the OV-associated cells were generated or whether their distribution reflects migration from the OV.

The ventricular surface also displayed a conserved molecular organization across species. Across the three species, enrichment of β-catenin and/or CD133 at the OV border was associated with markers of ciliary structures, including acetylated α-tubulin, ARL13B and AC3 (Berg et al., 2013; Dubreuil et al., 2007; Mirzadeh et al., 2008). The convergence of several markers is important because none is individually sufficient to demonstrate a specialized ciliated ventricular cell population. This is particularly evident for acetylated α-tubulin, which was strongly enriched at the sheep ventricular border but was also abundant in fibrillary structures within the parenchyma, consistent with its broader association with stable microtubules. In combination with the spatially restricted ARL13B and punctate AC3 signals, however, these observations support the maintenance of a differentiated apical/ciliary interface at the persistent adult OV. Such an interface may be biologically relevant because adult SVZ neural stem cells retain specialized apical domains contacting the ventricular lumen and bearing a primary cilium, providing a potential interface with signals present in the cerebrospinal fluid (Mirzadeh et al., 2008; Tong et al., 2014). Primary cilia also participate in signaling pathways involved in the regulation of specific neural progenitor populations, including Hedgehog signaling (Tong et al., 2014).

Despite pronounced differences in ventricular morphology, the persistent OV therefore shares several cellular features across species: progenitor- and glial-associated markers, proliferative cells, immature neuronal populations and a differentiated apical/ciliary interface. This suggests that lumen size or degree of patency alone may not determine whether the OV retains biologically relevant ventricular organization. Instead, preservation of the cellular architecture associated with the ventricular border may be the more informative feature.

The coexistence of these features is compatible with the hypothesis that the adult OV retains properties of a ventricular neurogenic compartment, but it does not yet establish an autonomous neurogenic niche. Even the identification of proliferating SOX2-positive cells at the sheep OV wall does not demonstrate that these cells generate the neighboring DCX-positive population. Likewise, the presence of proliferative and immature neuronal cells around the CD1 OV cannot exclude recruitment from the canonical SVZ-RMS system. Establishing local neurogenesis will therefore require direct birth-dating and lineage-tracing approaches capable of distinguishing OV-associated progenitors from cells originating in the SVZ. Such experiments will also be necessary to determine whether the cellular differences observed among species represent variations of a common neurogenic architecture or differences in the functional contribution of the OV to adult OB plasticity.

The convergence of evolutionary and cellular observations nevertheless changes the framework in which the adult OV can be considered. Rather than a uniformly transient ventricular extension, the OV appears to be an ancestral component of mammalian OB organization that persists into adulthood across much of mammalian diversity and retains a specialized cellular architecture in the species examined here. We therefore propose that, in mammals retaining an adult OV, this structure may represent an additional local progenitor compartment within the OB, spatially distinct from the canonical SVZ-RMS system. Whether OV-associated progenitors generate functional interneurons, whether they constitute a source of newborn neurons independent of the SVZ, and whether evolutionary variation in OV persistence contributes to differences in olfactory plasticity remain important questions for future investigation.

## 5. Conclusion

Our study challenges the conventional view of the OV as a transient developmental structure by showing that its persistence into adulthood is widespread across mammalian diversity and most likely represents an ancestral feature of the mammalian brain. Its absence is comparatively rare, phylogenetically structured and consistent with repeated secondary losses, while the intraspecific variation observed in Mus musculus and Rattus norvegicus further indicates that adult OV persistence can vary over relatively short evolutionary scales. Importantly, despite marked interspecific differences in OV morphology, comparative histological analyses reveal a conserved and organized ventricular architecture characterized by progenitor- and glial-associated markers, proliferative cells, immature neuronal populations, and a differentiated apical/ciliary interface. Although these observations do not establish local neuronal production or demonstrate that the OV constitutes an autonomous neurogenic niche, they show that the persistent adult OV is not simply an anatomically residual lumen but a specialized component of mammalian OB organization. Together, our findings provide an evolutionary and cellular framework for investigating the developmental mechanisms controlling OV persistence and for determining whether this ventricular compartment contributes to adult olfactory neurogenesis and plasticity.

## 7. Acknowledgements

The authors thank the MNHN imaging facility for access to the Zeiss LSM 880 Airyscan confocal microscope and for technical support with confocal imaging.

## 8. Conflict of Interest

The authors have no competing interests to declare

## 9. Author contributions

LB: Conceptualization, Methodology, Investigation, Formal analysis, Data curation, Visualization, Writing - original draft, Writing - review & editing, Project administration. J-PD: Resources. MM: Supervision, Writing - review & editing. SR: Supervision, Writing - review & editing.

## Notes

### Competing Interest Statement

The authors have declared no competing interest.

